# Transfected plasmids have reduced expression in cells deficient in SEPTIN 9 or ESCRT proteins

**DOI:** 10.64898/2026.08.21.746337

**Authors:** Emmanuel Ngwoke, Julie Hollien

## Abstract

Transfection of cells with DNA plasmids typically involves the uptake of lipoparticles by endocytosis, followed by the inefficient escape of these particles from endosomes into the cytoplasm. We found that the expression of transfected plasmids was reduced in cells depleted of either SEPTIN 9 or proteins in the endosomal sorting complexes required for transport (ESCRT) pathway. The reduction in plasmid expression could not be fully explained by effects on endocytosis. SEPTIN 9 depletion appeared to reduce the acidification of plasmid-containing compartments, suggesting that it primarily affects the pH-sensitive escape of plasmids from endosomes. Depletion of the ESCRT proteins VPS36 or ALIX resulted in especially dramatic reductions in transfected plasmid expression, which were accompanied by reduced colocalization between the transfected DNA and CHMP4, an ESCRT protein important for endosomal membrane remodeling during intraluminal vesicle formation. Finally, transfected plasmid DNA was strongly colocalized with LC3B, suggesting that the default pathway for transfected material is autophagy.

## Introduction

Lipid-based gene delivery systems offer advantages over viral-based methods, which tend to be more cytotoxic and prone to triggering immune responses (1). In lipofection, cationic liposomes are complexed with DNA to create lipoplexes, which attach to the plasma membrane and are taken up by endocytosis or pinocytosis. In order to avoid degradation in lysosomes, these lipoplexes must then escape from endosomal compartments into the cytosol. The DNA can then enter the nucleus and access transcription machinery.

Endosomal escape is considered the major bottleneck in the expression of DNA delivered by many types of lipid-based systems, and yet is perhaps the least-understood step in the process (2). Endosomes progressively acidify following their budding from the plasma membrane, and cationic lipoplexes have been designed to fuse with or disrupt the endosomal membrane as they become more acidic, leading to the release of the DNA. Despite advances in methods to detect the endosomal escape of transfected nucleic acids (3,4), we still lack a complete understanding of which aspects of endosomal maturation are important for this process.

Septins are small GTPases that play important roles in a variety of cellular processes, including cell division, endo-lysosomal trafficking, exocytosis, and cell migration (5,6). They can assemble into heterooligomeric filaments that bind to membrane surfaces as well as cytoskeletal structures, and are thought to function primarily as scaffolds. In mammalian cells, SEPTIN 9 (SEPT9) appears to also function independently of other septin family members in its ability to mediate interactions between endosomes/lysosomes and dynein motors, promoting the inward trafficking of these organelles on microtubules (7,8).

In addition to acidification and trafficking, endosome maturation is accompanied by the appearance of intraluminal vesicles, which form through the inward budding of the limiting membrane of the endosome, resulting in the encapsulation of membrane proteins targeted for degradation inside the organelle (9,10). The Endosomal Sorting Complexes Required for Transport (ESCRT) machinery coordinates this process through the sequential recruitment of four multimeric core complexes (ESCRT-0, -I, II, and -III) to the endosomal membrane (11). These complexes sort cargo proteins and carry out the membrane remodeling events that result in the budding of intraluminal vesicles. The endosomal maturation process culminates in the fusion of the resulting multivesicular bodies (MVBs, also known as late endosomes) with lysosomes and the degradation of the enclosed macromolecules. In addition to MVB biogenesis, the ESCRT machinery carries out several other membrane-remodeling events (11), including viral budding and cytokinesis.

Here we show that depletion of either SEPT9 or components of the ESCRT pathway leads to substantial reductions in the efficiency of transfected plasmid DNA expression, which we propose is a consequence of the reduced ability of endosomes to acidify and recruit membrane remodeling machinery, respectively.

## Materials and methods

### Plasmids, cell culture, and transfections

We obtained HTT-GFP plasmids from the Huntington Disease (HD) community biorepository (pCH00040 for HTT-Q145 and pCH00023 for HTT-Q73) and subcloned the coding sequences into an expression vector containing the EF1alpha promoter as previously described (13). We obtained CY3 and fluorescein labelled plasmids from Mirus Bio.

We cultured MC3T3-E1 cells (ATCC, RRID:CVCL_0409) in MEMα media containing nucleosides, L-glutamine, and no ascorbic acid (Life Technologies) with 10% Fetal Bovine Serum (FBS) at 37 C and 5% CO2. For siRNA-mediated silencing, we transfected cells with a mix of two siRNAs (Sigma-Aldrich) per target mRNA using the transfection reagent RNAi MAX (Invitrogen). As control, we used a commercial nontargeting siRNA from Sigma-Aldrich. We waited 48 h to allow cells to recover and degrade the targeted mRNA and protein before further transfection or analysis.

For transient DNA transfections, we transfected 0.5 μg of plasmid DNA except when testing the effects of increased DNA as noted in Fig 1F, using either Lipofectamine 2000 (Invitrogen) or polyethylenimine (PEI, from Polysciences), following the manufacturers’ protocols. We incubated the cells in 6 well plates with the transfection mixes at 37 C for 4 h (for Fig 1F) or 2 h (for all other experiments), then either collected cells or changed the media for later collections. Volumes were 2.0 for initial plating per well, 500 uL for the siRNA transfection mixes, and 250 uL for the plasmid transfection mixes, for a final volume per well of 2.75 mL.

**Fig 1.**
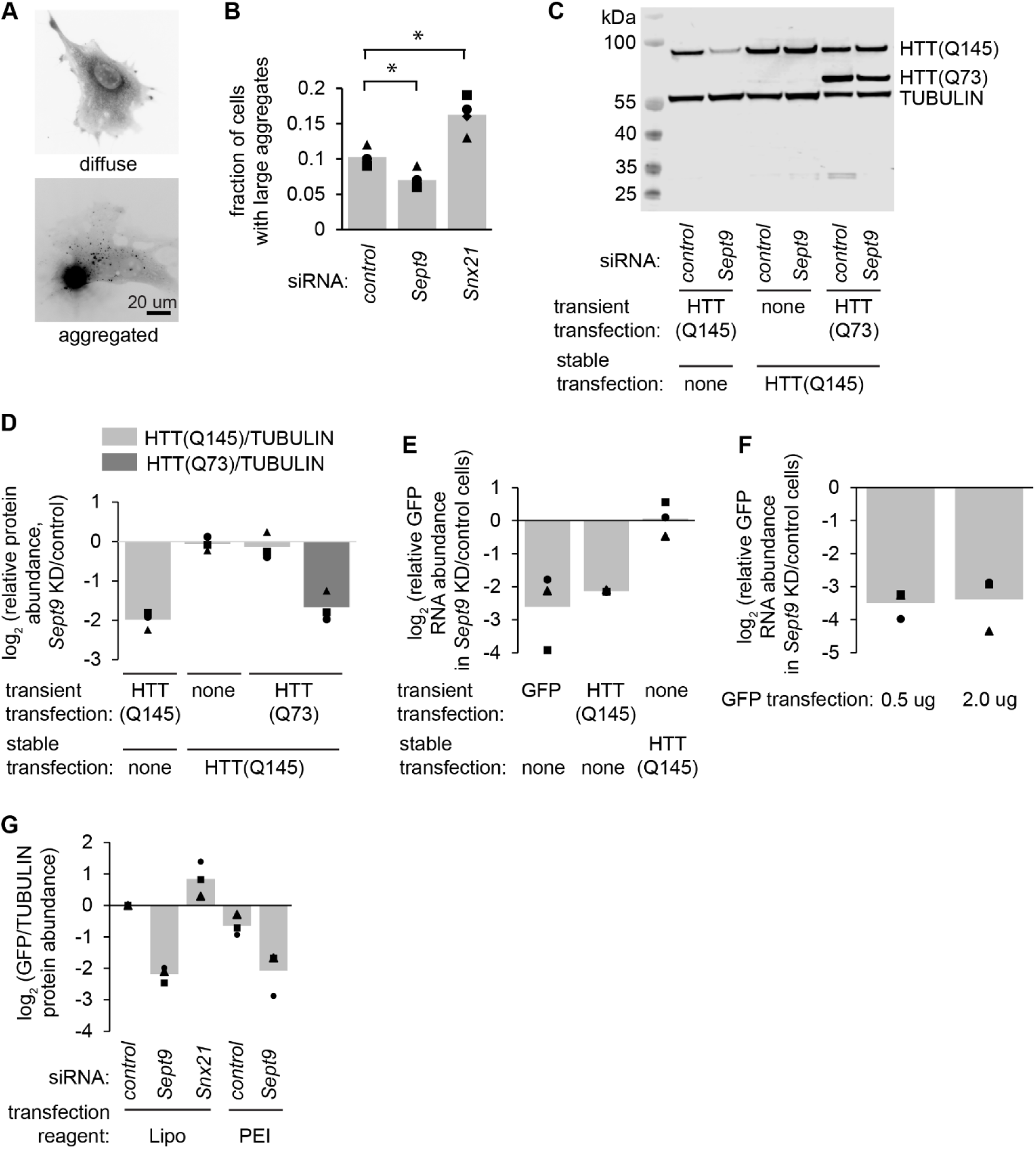
SEPT9 depletion reduces the expression of transfected plasmid DNA. (A) We transfected MC3T3-E1 cells with a plasmid encoding HTT-Q145-GFP and examined by fluorescence microscopy. Representative images show cells with diffuse vs aggregated HTT-Q145-GFP. (B) We depleted cells of the indicated mRNAs then transfected as in A and counted the fraction of transfected cells that displayed large juxtanuclear aggregates. Symbols indicate the average fractions for each of four independent experiments (100 cells per condition), and bars show the average of all experiments. *, p<0.05, paired t test followed by a Holm-Bonferroni correction for multiple pairwise comparisons. (C, D) We used siRNAs to deplete cells of *Sept9*, using a non-targeting siRNAs as a control, then transiently transfected plasmids expressing either HTT-Q145-GFP or HTT-Q73-GFP, into wild-type cells (lanes 1-2) or into cells stably expressing HTT-Q145-GFP (lanes 3-6). We then collected protein and analyzed by immunoblot using antibodies for GFP and TUBULIN (as a loading control). (D) shows the quantification of three independent replicates. (E, F) We depleted cells of *Sept9* as in C transfected plasmids encoding either GFP or HTT-Q145-GFP, then collected total RNA and analyzed by quantitative RT-PCR. (G) We depleted cells of the indicated mRNAs, then transfected a plasmid encoding GFP using either Lipofectamine 2000 (Lipo, as in all other figure panels) or polyethylenimine (PEI) and analyzed the GFP protein levels by immunoblot.

### Immunoblotting

For immunoblot analysis, we harvested protein by trypsinizing cells, washing with PBS, and lysing in RIPA buffer (25 mM Tris, pH 7.6, 150 mM NaCl, 1 % NP-40, 1% Na-deoxycholate, and 0.1% SDS) with protease inhibitors (Thermo Fisher Scientific, A32953). We determined protein concentrations using a Pierce BCA protein assay kit (Thermo Fisher Scientific, 23227) and a BSA (Thermo Fisher Scientific, 23210) standard curve. We resolved proteins using 4-12% polyacrylamide NuPage Bis—Tris gels. We transferred proteins to nitrocellulose membranes and incubated for 1 h in 5% dry milk diluted in TBS (20 mM Tris Base, 150 mM NaCl, pH7.4) with 0.02% Tween20. We prepared primary and secondary antibodies in 1% BSA, 0.1% Triton X-100 and 0.05% Tween20 in TBS. We then incubated the blots with anti-GFP (Invitrogen Molecular Probes, RRID: AB_221570, 1:7500) and anti-Tubulin (Cell Signaling Technology, RRID: AB_2210548, 1:5000) primary antibodies at 4 C overnight, washed, then incubated with the secondary antibody (IRDye 800CW Goat anti-Rabbit IgG, Licor, RRID: AB_621843, 1:10,000, 1h). We scanned the membranes using a LiCor Odyssey CLx Imager and quantified band intensities using the LiCor Image Studio Software.

### Quantitative PCR

We extracted and DNase-treated RNA using the Quick-RNA Miniprep Kit (Zymo Research), which incorporates an on-column DNase I treatment step during RNA purification, with an incubation period of 20 minutes, to eliminate genomic DNA contamination. We additionally included a reverse transcription-negative control (RT-) to confirm the absence of residual genomic DNA. We assessed RNA concentration and purity using a NanoDrop 1000 spectrophotometer (Thermo Fisher Scientific). All samples had 260/280 ratios between 2.06–2.11 and 260/230 ratios between 1.91–2.27. We used 1 µg of RNA as the input for reverse transcription. We synthesized cDNA using Moloney Murine Leukemia Virus Reverse Transcriptase (MMLV-RT, New England Biolabs) with an oligo-T18 primer. We performed qPCR on a QuantStudio 3 real-time PCR system (Life Technologies) in a total reaction volume of 25 µL with a heated cover temperature of 105°C, using SYBR Green as the fluorescent detection dye and a primer concentration of 50 µM. Cycling conditions were as follows: initial denaturation at 95°C for 2 minutes followed by 40 cycles of denaturation at 95°C for 20 seconds, annealing at 55°C for 20 seconds, and extension at 68°C for 40 seconds. We performed a melt curve analysis after amplification to confirm amplicon specificity, consisting of 95°C for 15 seconds, 60°C for 15 seconds, a continuous dissociation step rising to 95°C at 0.1°C/s, and a final hold at 25°C for 1 minute. We measured each sample in triplicate and quantified relative RNA abundance for Gfp using Rpl19 as a housekeeping reference gene against a serially diluted standard curve. We selected Rpl19 as the reference gene as it has been widely validated as a stable MC3T3-E1 housekeeping gene under similar experimental conditions. We analyzed data using the Thermo Fisher cloud-based platform (app.thermofisher.com). Standard curve analysis yielded R² values that were over 0.980. The following primer sequences were used: GFP forward: 5’ TCATCTGCACCACCGGCAAG, GFP reverse: 5’ CAGCTCGATGCGGTTCACCA (GFP amplicon size = 219 nucleotides), Rpl19 forward: 5’ CTGATCAAGGATGGGCTGAT, Rpl19 reverse: 5’ GCCGCTATGTACAGACACGA (Rpl19 amplicon size = 490).

### Flow cytometry

We trypsinized and washed cells, resuspended in 150 μl PBS, and analyzed 10,000 cells per condition using a Beckman Coulter Cytoflex S instrument, gating for live, singlet cells. We determined the fraction of transfected cells by comparing to mock-transfected control cells and calculated the median intensity of all cells for each condition using the CytExpert software v2.6 (Beckman Coulter Cytoflex S).

### Immunostaining and microscopy

For imaging HTT-GFP in live cells, we grew MC3T3-E1 cells in glass-bottom dishes and carried out transfections as described above. For immunostaining, we grew MC3T3-E1 cells on glass coverslips, and transfected with 0.5 μg of CY3-labelled plasmid. We made all fixation and staining solutions in Phosphate-Buffered Saline containing 1 mM MgCl. To improve the background cytosolic signal in the CHMP4 staining, we gently permeabilized the cells prior to fixing using digitonin (10 μg/ml, 4 C, 15 min). This step was not necessary for LC3 staining. We fixed cells with 4% paraformaldehyde (37 C, 15 min) and permeabilized with 0.2% Triton-X (room temperature, 20 min). We then blocked with 2% BSA and 0.02% tween-20 (room temperature, 10 min) and stained with CHMP4B (1:300, Proteintech, RRID: AB_2877971) or LC3 (1:250, Proteintech, RRID: AB_2137737) primary antibodies. We washed the coverslips with 0.02% tween-20 and stained with a secondary antibody (Alexa-Flour 488 anti-rabbit IgG, Invitrogen A11008, RRID: AB_143165, 1:1000), and washed again before mounting the coverslips using ProLong Diamond Antifade Mountant with DAPI (Invitrogen).

We imaged the cells at room temperature using an Olympus IX-51 inverted epifuorescence microscope with a 60X (NA 1.25) oil objective, an Olympus DP23 Monochrome camera, and the CellSens Standard v3 acquisition software (typical resolution limit ∼0.22 µm). The light source was a CoolLED pE-300White Direct Couple Single Band / UV, set to 50% power. Exposure times were: 500 msec (HTT-GFP), 50 msec (Cy3), 300 msec (Chmp4 and LC3).

### Statistics

All replicates shown in the figures are biological (not technical) replicates carried out on different days with different populations of cells. For most comparisons we used Student’s t-tests to carry out pairwise statistical tests, and corrected for multiple comparisons using the Holm-Bonferroni method. For single tests we used a significance cutoff of p<0.05. For the Holm-Bonferroni corrections we multiplied the lowest p-value by the number of tests (n), and the second lowest p-value by (n-1), etc until a cutoff of p >= 0.05 was reached. For datasets with more than 8 comparisons (ie, Fig 3A) we instead used a one-way ANOVA followed by a Tukey’s Honest Significant Difference with a significance cutoff of p<0.05.

## Results and discussion

### SEPT9 depletion impairs the expression of transfected plasmid DNA

We began this study by investigating the role of SEPT9 in the aggregation of the Huntingtin protein, which underlies Huntington’s disease (12). Pathological versions of this protein contain long stretches of glutamine residues that promote its aggregation in distinct juxtanuclear structures, and our previous work suggested that trafficking of endosomes and lysosomes to this same area of the cell reduces the accumulation of aggregates (13). We therefore hypothesized that depletion of SEPT9, which would reduce this inward trafficking of lysosomes, would lead to increased Huntingtin aggregation. To test this, we depleted cultured mouse MC3T3-E1 cells of SEPT9 via RNA interference (RNAi), then transfected with a plasmid expressing an N-terminal fragment of the Huntingtin protein with a stretch of 145 glutamine residues tagged with GFP (HTT-Q145-GFP), using the transfection reagent Lipofectamine 2000. As expected, cells expressing HTT-Q145-GFP displayed either diffuse GFP fluorescence throughout the cytoplasm or a large juxtanuclear aggregate (Fig 1 A). These aggregates were distinctive, bright, next to the nucleus, and approximately 5-10 um in diameter (Fig 1 A). Although we occasionally saw smaller aggregates distributed through the cytoplasm, we did not include these in the analysis. As a positive control, depleting cells of sorting nexin 21 (SNX21), which recruits Huntingtin to endosomes and thereby assists in its degradation (14), increased the fraction of cells containing these juxtanuclear aggregates (Fig 1 B). Contrary to our hypothesis, depleting cells of SEPT9 significantly reduced the fraction of cells containing aggregates (Fig 1 B).

The reduced aggregate formation in SEPT9-depleted cells could be explained by reduced HTT-Q145-GFP protein expression, which we confirmed by immunoblot (Fig 1 C). The reduced aggregate formation in SEPT9-depleted cells could be explained by reduced HTT-Q145-GFP protein expression, as seen by immunoblot (Fig 1 C). To test whether these reduced protein levels were a consequence of reduced transfection efficiency, we generated a cell line in which the same HTT-Q145-GFP plasmid was stably integrated into the genome. In these stably transfected cells, SEPT9 depletion had no effect on HTT-Q145-GFP protein levels (Fig 1 C and D). To test whether SEPT9 affected expression from transiently transfected plasmids only or if the transfection process indirectly led to reduced expression of HTT-Q145-GFP, we transiently transfected the stable cell line with a related plasmid expressing HTT-GFP with a shorter polyglutamine repeat (HTT-Q73-GFP). In this experiment, SEPT9 depletion led to reduced protein levels for the transiently transfected HTT-Q73-GFP, while not affecting the protein levels of the stably expressed HTT-Q145-GFP (Fig 1 C,D).

These results suggested that SEPT9 depletion specifically diminishes expression of transiently transfected plasmids. To confirm that this effect was caused by reduced levels of the mRNA generated from the transfected DNA, we repeated the RNAi and transfection experiments and collected RNA for analysis by real-time quantitative PCR (RT-PCR). Similar to the results from the protein analysis, SEPT9 depletion led to a reduction in HTT-Q145-GFP mRNA levels in transiently transfected cells but not in cells stably expressing the same mRNA (Fig 1 E). This reduction was not specific to HTT-GFP, as we saw a similar reduction in mRNA levels for transiently transfected GFP (Fig 1 E). Furthermore, this was not dependent on the amount of plasmid DNA transfected, as comparing 0.5 µg to 2.0 µg yielded identical effects (Fig 1 F).

We next asked if the effect of SEPT9 depletion was specific to our Lipofectamine 2000 transfection reagent. We repeated the SEPT9 RNAi experiment using a non-lipid polyethylenimine (PEI) reagent to transfect a plasmid expressing GFP. Western blot analysis showed a ∼4-fold decrease in GFP protein levels in SEPT9-depleted cells (Fig 1 G). Taken together, these results show that depleting cells of SEPT9 impairs the expression of transiently transfected plasmids in a manner that is independent of the transfection reagent.

### SEPT9 depletion slightly reduces the uptake of labeled plasmids

Expression of transiently transfected plasmids could be impaired by inhibiting the endocytic uptake of lipoplexes, the escape of plasmid DNA from endocytic organelles into the cytosol, the entry of plasmids into the nucleus, and/or the downstream processes of transcription. To test whether the first step in this process was affected by SEPT9, we transiently transfected cells with plasmid DNA fluorescently labeled with CY3 and measured the plasmid uptake by flow cytometry. After 2 h of incubation with lipofectamine-plasmid complexes, approximately 80% of cells showed CY3 fluorescence above the background levels of mock-transfected cells, and this transfection efficiency was not affected by SEPT9 depletion (Fig 2 A). SEPT9 depletion did reduce the median fluorescence by ∼20% (Fig 2 B), suggesting a modest decrease in the amount of material taken up by each cell, though this difference was not statistically significant and not on a scale that would explain the ∼4-fold decrease in mRNA levels (Fig 1 E and F).

**Fig 2.**
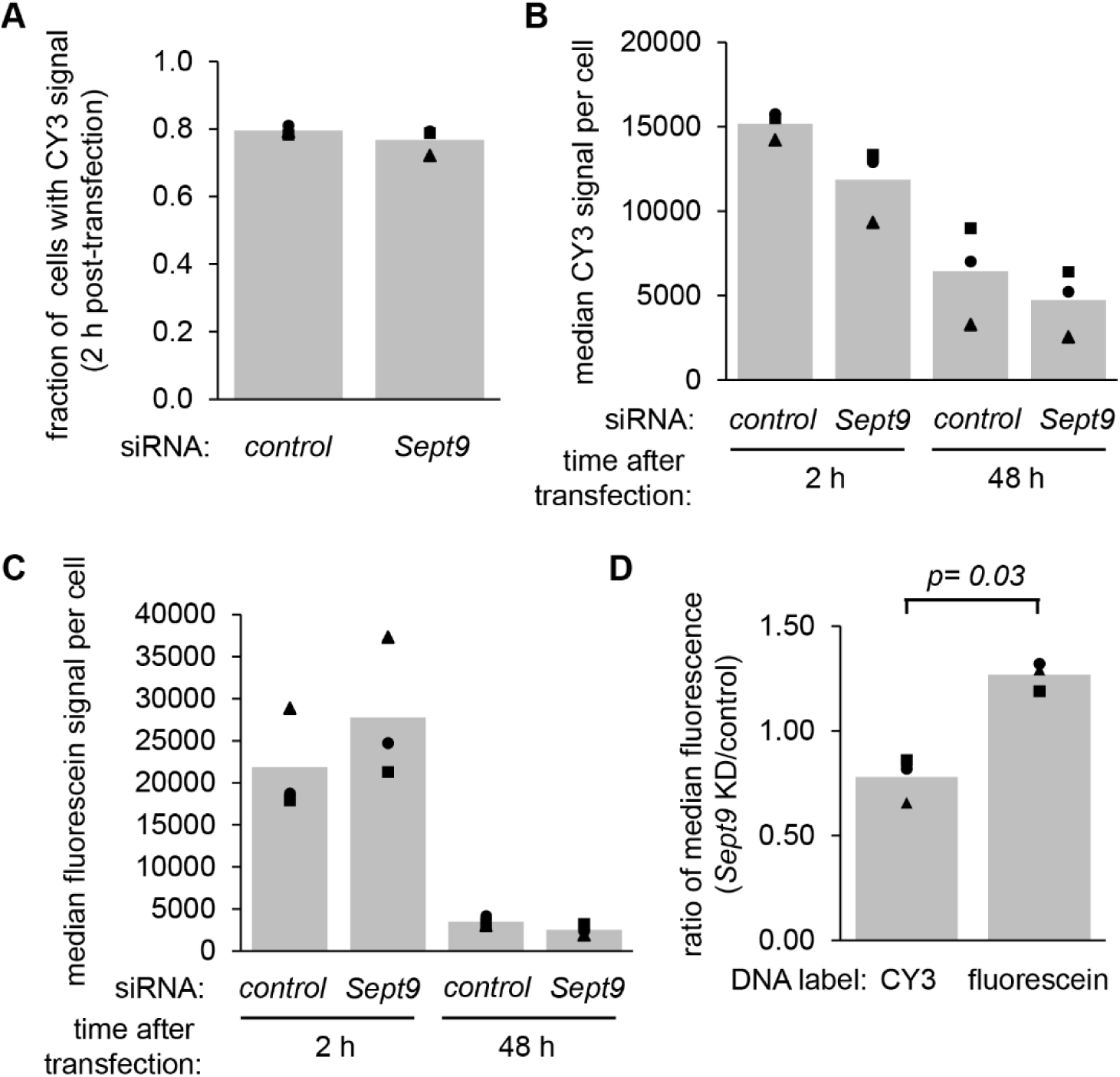
SEPT9 depletion slightly reduces the uptake of labeled plasmids but results in less quenching of fluorescein-labeled transfected DNA. (A,B) We used siRNAs to deplete MC3T3-E1 cells of *Sept9*, using a non-targeting siRNAs as a control, then transfected with a CY3-labeled plasmid. After incubating cells with the transfection lipoplexes for 2 h, we either harvested cells immediately or changed the media and harvested after 48 h. We then analyzed by flow cytometry. (A) shows the fraction of cells with CY3 fluorescence above background levels, and (B) shows the median CY3 signal per cell for 3 independent experiments (bars show the average of the experiments). (C) We repeated the experiment in A,B using a plasmid labeled with the pH-sensitive fluorescein. (D) shows the ratio of fluorescence signals in *Sept9* depleted vs control cells, following 2 h incubation with transfection lipoplexes containing the CY3 or fluorescein-labeled plasmids.

In contrast to the CY3-labeled plasmid results, transfection with fluorescein-labeled plasmids showed an increase in median fluorescence of ∼27% for the SEPT9-depleted cells (Fig 2 C and D). Although the data did not pass a Student’s t-test for comparing the median fluorescence signals in control vs. SEPT9-depleted cells (p = 0.06), the ratios between controls and knockdown cells for the CY3 vs. fluorescein-labelled DNA transfections were statistically different. This difference could be explained by the fact that unlike CY3, fluorescein is pH sensitive and quenched in acidic environments such as late endosomes and lysosomes (15). Consistently, the median fluorescein signal diminished to near-baseline levels after 48 h, when most plasmid DNA would be expected to have reached the acidic lysosomes or be degraded.

These results suggest that SEPT9 depletion has only mild effects on the uptake of plasmids during lipofection but potentially impairs their expression by slowing or reducing the acidification of endosomes containing the entrapped plasmids, preventing their release into the cytosol and ultimately their nuclear delivery. This effect on endosomal escape is consistent with a recent study showing that certain endosomes bind preferentially to septin-coated microtubules, slowing their trafficking and promoting their maturation (16). The maturation steps impeded by depleting cells of septins include acidification and the acquisition of lysobisphosphatidic acid (LBPA), a negatively-charged phospholipid that is not only important for the formation of intraluminal vesicles but is also thought to be critical for the ability of lipoplexes to permeabilize the endosomal membrane (17).

### ESCRT proteins affect the expression of transfected plasmid DNA

To examine the effects of other endosomal factors in the expression of transfected plasmids, we depleted cells of a series of ESCRT proteins (Fig 3 A), transfected a GFP-expressing plasmid, and monitored the expression of the GFP mRNA by RT-PCR. Each of these siRNA-mediated knockdowns reduced the expression of the transfected plasmid, compared to non-targeting siRNAs or the additional control of SNX21. Depletion of VPS36 or ALIX had the most dramatic effects, reducing mRNA levels by over 50-fold (Fig 3 A), surpassing the effects of SEPT9 depletion by an order of magnitude.

**Fig 3.**
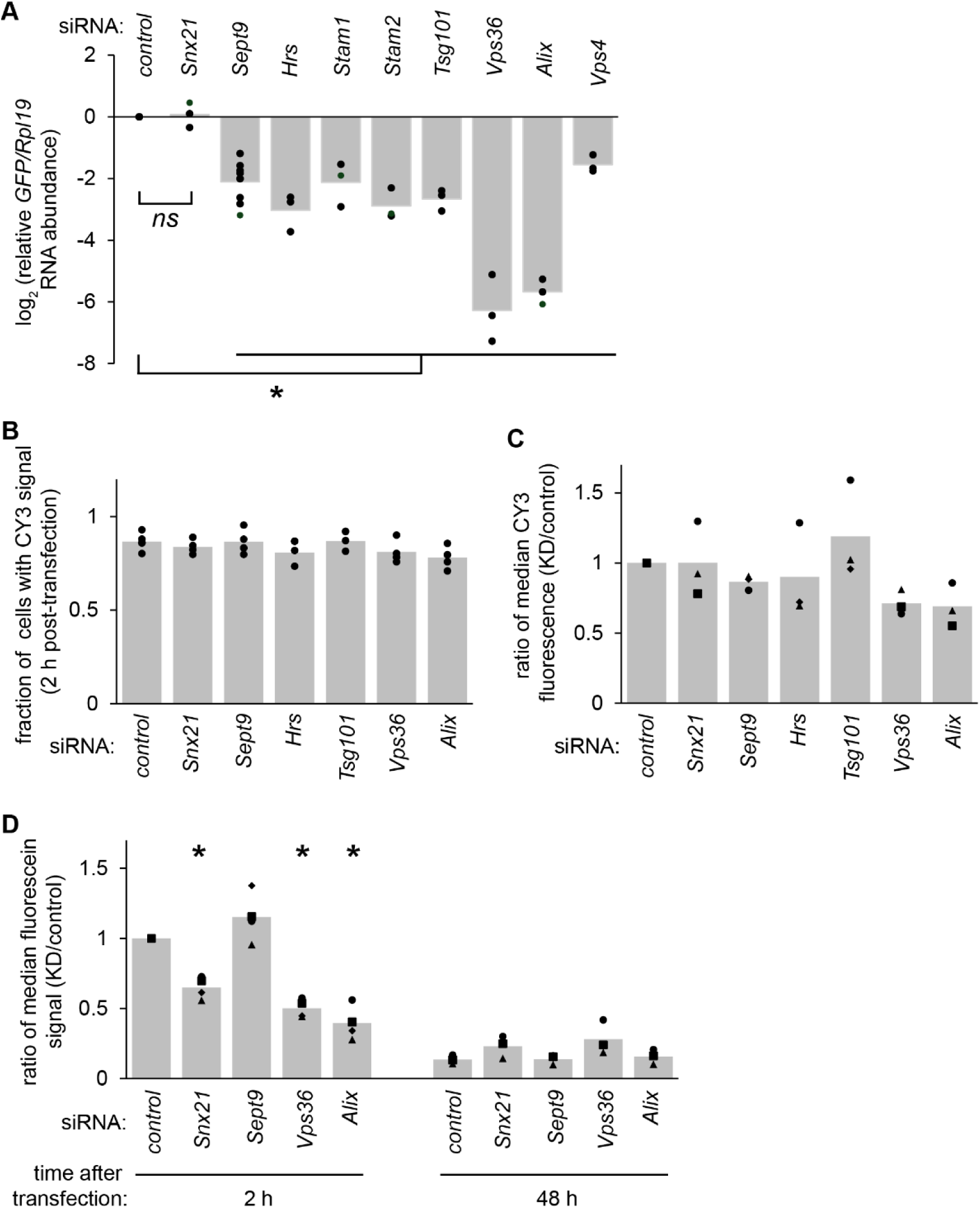
ESCRT depletion reduces the expression of transfected plasmid DNA. (A) We used siRNAs to deplete MC3T3-E1 cells of the indicated factors, transfected a plasmid encoding GFP, collected RNA after 48 h, and analyzed by quantitative RT-PCR. Symbols represent the relative RNA level in each independent experiment and bars show the average of all experiments. (B,C) We depleted cells of the indicated factors, then transfected with a CY3-labeled plasmid and analyzed by flow cytometry. (B) shows the fraction of cells with CY3 fluorescence above background levels, and (C) shows the median CY3 signal per cell relative to the control knockdowns. (D) We repeated the experiment in B,C using a plasmid labeled with the pH-sensitive fluorescein.

Similar to SEPT9 depletion, ESCRT depletion did not appear to affect the fraction of cells that were transfected with the CY3-labeled plasmid (Fig 3 B), and affected the median CY3 fluorescence per cell by less than 2-fold (Fig 3 C). Unlike SEPT9 depletion, VPS36 and ALIX depletion also reduced the fluorescence of transfected fluorescein-labeled plasmids to a similar degree as seen for CY3 (Fig 3 D). These results suggest that the primary effect of ESCRT depletion was mediated by neither the initial uptake of plasmids into the cell nor the acidification of the endosomal structures containing the plasmids.

### VPS36 or ALIX depletion reduced CHMP4 recruitment to endosomes containing plasmid DNA

VPS36 is a component of ESCRTII, whose main function is to recruit the proteins that comprise ESCRTIII, including the key factor CHMP4B (11). ESCRTIII proteins polymerize on the cytoplasmic face of endosomal membranes to carry out the membrane remodeling and scission events that result in inward budding and detachment of vesicles from the endosomal membrane (18,19). ALIX is also thought to help position ESCRTIII on the membrane and bridge CHMP4 oligomers (20). Notably, our attempts to deplete cells of CHMP4B resulted in substantial cell death, preventing us from testing the effects of ESCRTIII on transfection efficiency.

Given that recruitment and organization of ESCRTIII is a common function for the two factors with the most dramatic effects on transfected plasmid expression, we sought to test whether CHMP4B localization to plasmid-containing endosomes was impaired in cells depleted of these factors. We depleted cells of SEPT9, VPS36, or ALIX, and incubated with CY3-labeled plasmid lipofectamine complexes for 2 h. We then fixed cells and stained using an antibody for CHMP4B. Most cells displayed CY3 fluorescence in discrete cytoplasmic foci, with an average of 3-4 foci per cell that was not affected by the knockdowns (Fig 4 A and B). The most notable phenotype in control cells was the presence of bright CHMP4B foci that were larger than typical endosomes and colocalized with CY3. We counted the fraction of cells that contained at least one of these structures, identified by their size of over 1 μm and their colocalization with both CHMP4 and CY3. Depletion of either VPS36 or ALIX, but not SEPT9, reduced the fraction of cells containing these structures, suggesting an impaired recruitment of CHMP4B to endosomes containing transfected plasmid DNA (Fig 4 C). Unfortunately, though not surprisingly given the low efficiency of plasmid endosomal escape, we did not detect any CY3 fluorescence in the cytosol or nucleus.

**Fig 4.**
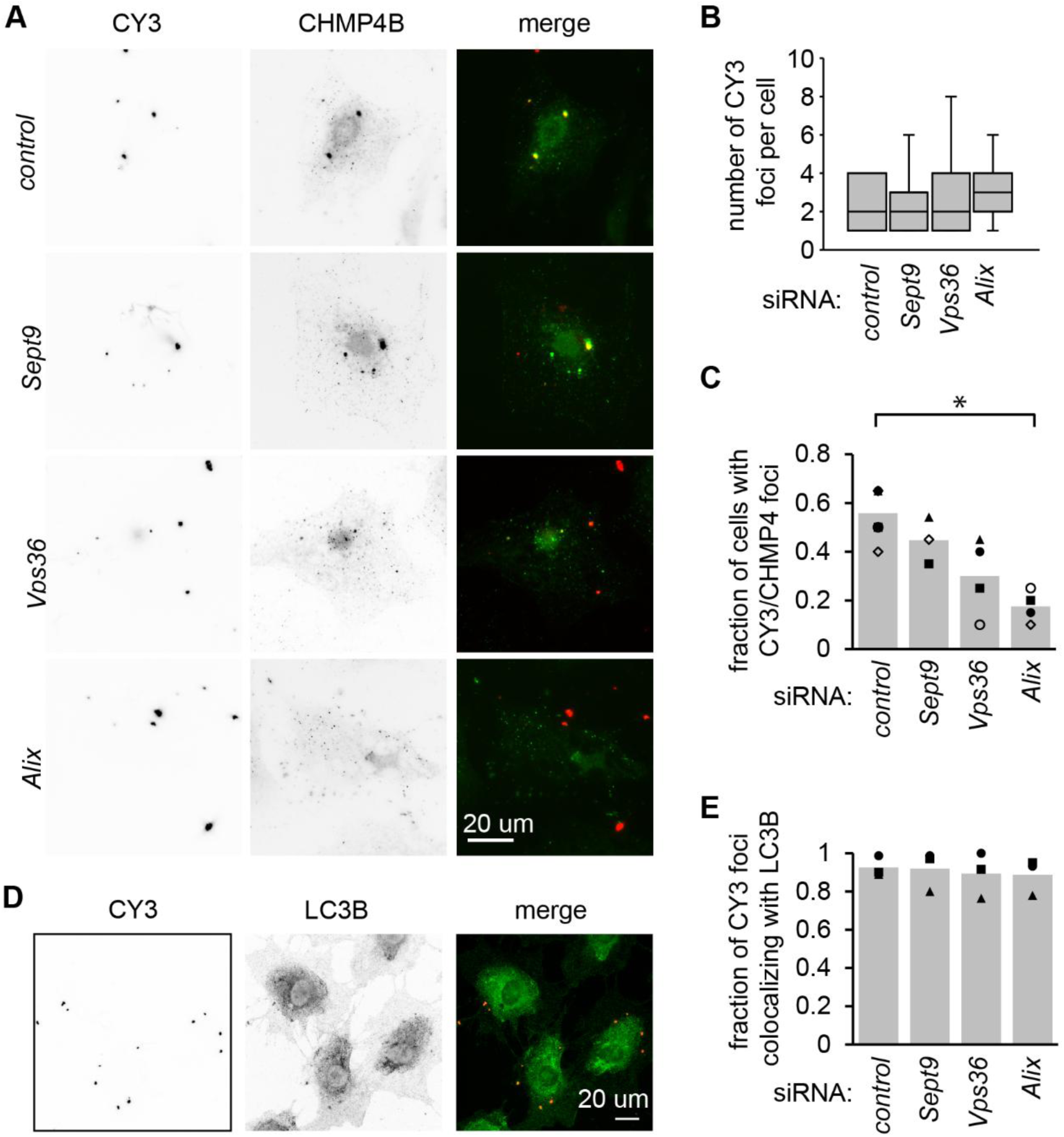
VPS36 or ALIX depletion reduces CHMP4 colocalization with plasmid DNA. (A-C) We used siRNAs to deplete MC3T3-E1 cells of the indicated factors, then transfected a CY3-labeled plasmid and fixed and stained cells for CHMP4. (A) shows representative images of the CY3 and CHMP4 signal and (B) shows the number of CY3 foci per cell (30 cells per experiment, 3 experiments; shown is a box plot of all 90 cells per condition). (C) shows the fraction of transfected cells containing at least one CHMP4 structure with a diameter of greater than 1 μm and colocalized with CY3. *, p<0.05, paired t test followed by a Holm-Bonferroni correction for multiple pairwise comparisons. (D, E) We repeated the experiment in A-C but stained for LC3B rather than CHMP4. (D) shows representative images from control cells, and (E) shows the fraction of CY3 foci that colocalized with LC3 (75-100 foci counted per experiment, 3 independent experiments).

### Transfected plasmids colocalize with the autophagy factor LC3B

We detected many CY3 foci that did not overlap with CHMP4B signal, leading us to question the identity of these structures. We repeated the siRNA treatments and CY3-plasmid transfections, this time staining for the autophagosome marker LC3B (Fig 4 D and E). Strikingly, almost every CY3 structure in all conditions colocalized with LC3 foci, suggesting that the endosomes containing transfected plasmids were targeted to autophagy.

### Perspective

The results described here suggest that transfected plasmids rely not only on the acidification of endosomes to accomplish their escape into the cytosol, but also on ESCRT machinery. ESCRTs may aid in endosomal escape by deforming the membrane as they promote intraluminal vesicle formation, resulting in weak points that the lipoplexes exploit. Alternatively, cells with deficient ESCRT machinery may target endosomes to autophagy more efficiently, either directly due to their abnormal ESCRT complement at the membrane (21) or by the failure of ESCRTs to repair endosomal membrane damage (22,23). This may in turn reduce transfection efficiency as lipoplexes become more quickly entrapped by autophagosomes, as previously described (3,24). Overall, our results suggest that manipulation of ESCRT-mediated membrane remodeling may be an effective approach for enhancing lipid-based gene delivery in mammalian cells.

## Acknowledgments

We thank the Hollien lab for helpful discussions and advice.

## Notes

### Competing Interest Statement

The authors have declared no competing interest.

### Funding

- National Institutes of General Medical Sciences: 2R35GM119540

